# A switching cost for motor planning

**DOI:** 10.1101/047621

**Authors:** Jean-Jacques Orban de Xivry, Philippe Lefèvre

**Affiliations:** KU Leuven, Department of Kinesiology, Movement Control and Neuroplasticity Research Group, Tervuursevest 101, 3001 Leuven, Belgium; Institute of Information and Communication Technologies, Electronics and Applied Mathematics and Institute of Neuroscience, Université catholique de Louvain, 1348 Louvain-La-Neuve, Belgium

## Abstract

Movement planning consists of choosing the endpoint of the movement and selecting the motor program that will bring the effector on the endpoint. It is widely accepted that movement endpoint is updated on a trial-by-trial basis with respect to the observed errors and that the motor program for a given movement follows the rules of optimal feedback control. Here, we show clear limitations of these predictions because of the existence of a switching cost for motor planning. First, this cost prevented participants from tuning their motor program appropriately for each individual trial. This was true even when the participants selected the width of the target that they reached toward or when they had learned the appropriate motor program previously. These data are compatible with the existence of a switching cost such as those found in cognitive studies. Interestingly, this cost of switching shares many features of costs reported in cognitive task switching experiments and, when tested in the same participants, was correlated with it. Second, we found that randomly changing the width of a target over the course of a reaching experiment prevents the motor system from updating the endpoint of movements on the basis of the performance on the previous trial if the width of the target has changed. These results provide new insights into the process of motor planning and how it relates to optimal control theory and to a selection by consequences process rather than to an error-based process for action selection.

## Introduction

Movement planning, which includes all the processes occurring before the movement (detection and selection of target, selection of feedback control policy, etc.) is widely thought to follow two distinct rules. First, the control policy (how the brain will react with respect to incoming sensory input during the movement) is tuned according to optimal control laws (Todorov and Jordan, 2002; Shadmehr and Krakauer, 2008; Diedrichsen et al., 2009; Pruszynski and Scott, 2012). Second, movement performance is adjusted on a trial-by-trial basis via learning from errors even when learning or adaptation are not explicitly required (van Beers, 2009; Verstynen and Sabes, 2011; Dingwell et al., 2013; Wong and Shelhamer, 2014). In this paper, we demonstrate that these two rules are violated as soon as the target width changes randomly over the course of trials.

Such disruption of performance due to randomization is widespread in motor tasks (Elliott and Allard, 1985; Edin et al., 1992; Horak and Diener, 1994; Khan et al., 2002; Pruszynski et al., 2008; Selen et al., 2009; Bennett et al., 2010; Afsanepurak et al., 2012) (but see White and Diedrichsen, 2013). For instance, in a recent study (Orban de Xivry, 2013), we asked participants to reach to either a narrow or wide target and found that participants reacted differently to a perturbation in function of the target width (compatible with the minimum intervention principle of optimal control, Todorov and Jordan, 2002) but unexpectedly also in function of the schedule (random or blocked). That is, the modulation of the response to perturbations with target width was smaller in the random (target width randomly changes from trial-to-trial) than in the blocked schedule.

Interestingly, decrease in performance due to randomization is also observed in cognitive tasks where it falls in the category of switching costs under the name of mixing cost. A switching cost relates to the inability of the participants to switch from one task to another without a penalty in behavior (Monsell, 2003; Kiesel et al., 2010). Typically, in cognitive studies, this cost is reflected in an increase in reaction time (Allport et al., 1994; Rogers and Monsell, 1995). For instance, when participants are required to report the color or the shape of a symbol based on some abstract rules, the reaction time is shorter when the rule is presented in blocked schedule (i.e. the rule remains unchanged for several trials in a row) than when the rule can change randomly from trial-to-trial. Interestingly, such switching cost is also present when participants choose the rule that they want to apply on each trial (Arrington and Logan, 2004, 2005).

While in the previous paper (Orban de Xivry, 2013), we merely reported the observation that randomly changing the width of the target affected the optimality of motor behavior, we present in this paper three experiments that provide an explanation for the absence of optimality in the random schedule and identify limits on the flexibility of motor planning. Importantly, in this study, like in many others (Ahmadi-Pajouh et al., 2012; Nashed et al., 2012; Dimitriou et al., 2013; Orban de Xivry, 2013; White and Diedrichsen, 2013), we consider that motor planning (i.e. before movement onset) consists in the derivation of a goal-directed feedback control law (Todorov and Jordan, 2002; Scott, 2004; Todorov, 2004; Liu and Todorov, 2007) and that responses to any perturbations during the movement are driven by this feedback law. That is, a perturbation at any time during movement allows us to probe the feedback control policy determined before movement (Liu and Todorov, 2007; Nashed et al., 2012; Gallivan et al., 2016). These perturbations reveal that humans cannot switch between two different control policies for a single trial in order to behave optimally even when they choose on each trial the width of the target that they are reaching to or when they have exhibited the optimal behavior in a different context. In addition, we found that participants who were better at switching between two instructions during a cognitive switching task were also better at switching between two motor control policies. Together, these studies provide new insights into the mechanisms of motor planning (Cisek and Kalaska, 2010; Wong et al., 2015).

## Methods

### Participants

Sixty-one healthy participants (21 for experiment 1 and 20 for experiment 2 and for experiment 3) were enrolled for the experiments after written informed consent. All participants had no history of neurological disorders, were right-handed and between 18 to 40 years old. All procedures were approved by the Ethics Committee of the Université catholique de Louvain. We also analyzed some new aspects of the data from the 50 participants from our previous paper (Orban de Xivry, 2013).

### Setup

Participants were sitting in front of a robotic arm (Endpoint Kinarm, BKin Technologies, Kingston, Ontario, Canada) that controlled a cursor that appeared to be positioned at the same position in space as the hand through a mirror setting. Participants could not see their hand and the displayed cursor was the only available visual feedback of their hand position. Hand position, velocity and acceleration and the force exerted by the participants on the handle of the robot were recorded at 1000Hz and stored on a computer for offline analysis. The robot was also able to exert forces in order to perturb or direct the hand of the participants.

### Protocol

#### Cognitive task switching

In this cognitive task switching, participants had to indicate either the color or the shape (depending on a specific instruction) of a symbol by applying a force with their right or left hand (Fig. 1, cognitive task switching). Depending on the schedule, the instruction varied randomly from trial-to-trial or remained unchanged for several trials in a row. More specifically, participants had to respond to the appearance of a visual stimulus (disk (radius: 1cm, surface: 3.14cm^2^) or square (1.75×1.75cm, surface: 3.06cm^2^) that was either blue or orange (these colors were colorblind-friendly) on the screen of the Kinarm setup by applying an isometric force of 7N on the left or right arm of the robot (bimanual task) following one of two possible rules. The direction of the force was irrelevant, only its magnitude was taken into account. For the shape rule, participants had to indicate whether the shape was a square or a circle. The association between shape and hand of the response was counterbalanced across participants. For the color rule, the participants had to indicate whether the symbol was orange or blue. The association between color and hand of response was counterbalanced across participants. On each trial, the relevant rule was indicated by a solid white line (20cm wide) that was displayed 1.5cm above the symbol for the shape rule and 1.5cm below the symbol for the color rule. When a participant made a mistake, a red line (20cm wide) appeared on top of the symbol. Intertrial time interval was 400ms. Participants had maximum 5s to provide their response.

**Figure 1.**
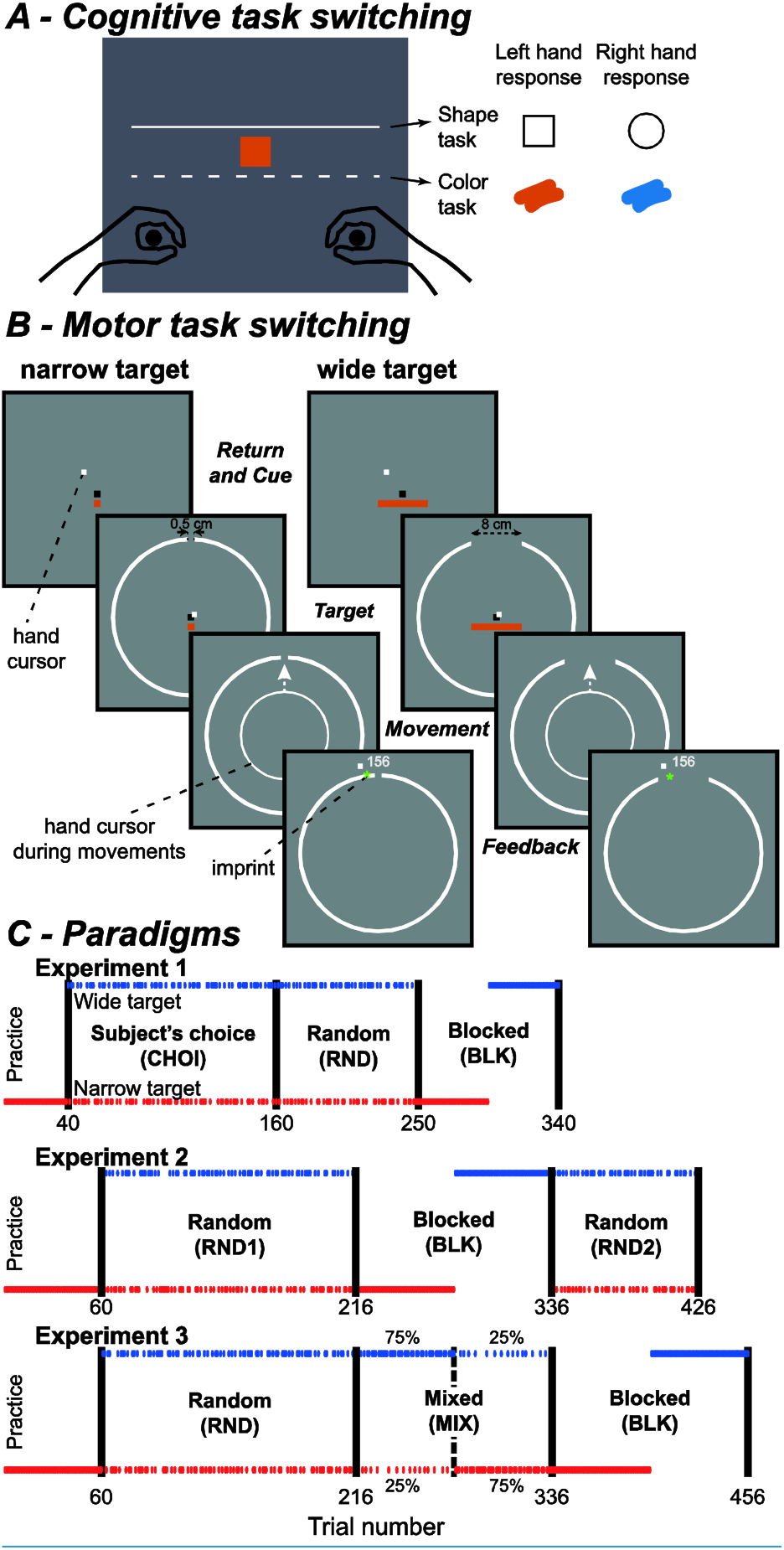
Experimental protocols. A) Cognitive task switching: participants had to indicate the shape of the symbol (shape task: square or circle) if a white line was displayed above the symbol on the screen (as on the panel) or its color (color task: orange or blue) if the line was below it (at the place of the dashed line). The response was indicated by exerting an isometric force on the left or right handle of the robot manipulandum. The situation depicted on the figure requires a left hand response. The hands were not visible for the participants. B) Motor task switching: In this task, participants were instructed to bring the cursor through the aperture of a circle. This aperture was always at the same location but could have one of two possible widths (0.8 or 8cm). Each trial started with the appearance of the starting position (Black square, return and cue phase). During this phase, a cue indicated the width of the aperture for the ensuing trial (orange symbol, Return and Cue phase). The target was then presented 15 cm away from the starting position in front of the participant (Target phase). When the participants initiated their reach, the hand cursor was replaced by a circle whose radius increased with the distance travelled by the hand (Movement phase). Veridical hand position was provided after movement end in order to provide feedback about movement accuracy (feedback phase). Additional information was provided via the imprint that the hand left on the circle when it crossed it. The imprint was blue if movement duration was too long, yellow if movement duration was too short and green if movement duration was between 500 and 600ms. One point could be earned for good movement speed and another point could be earned for good accuracy. The total number of points collected so far during the block was displayed at the end of each trial (156 points in the illustration). C) The three panels illustrate the repartition of trials with the narrow and wide target over the course of the three experiments (see main text). The y-axis represents target width and the x-axis trial number. In the CHOI schedule, participants selected the width of the target for each trial and were instructed to do so randomly. In the random schedule, target width was randomly varied from trial-to-trial. In the blocked schedule, target width remained unchanged for either 45 trials (experiment 1) or 60 trials (experiments 2 and 3). The order of target width was counterbalanced across participants. In the MIX schedule, there was an imbalance in the percentage of trials for each target width (75% - high probability trials vs 25% - low probability trials). Each participant received 60 trials with higher probability of one target width followed by another series of 6o trials where the other target width was more frequent. The order was also counterbalanced across participants.

After one to three blocks of training to demonstrate the task (20 trials each), the participants performed three blocks of trials. In the random schedule (first and third blocks, 40 and 64 trials), the rule (indicate shape or indicate color) could change pseudorandomly from trial to trial. In contrast, during the second block, the rule remained unchanged for the first half of the block (20 trials) after which it switched to the other rule until the end of the block (20 trials).

#### Motor task switching

Participants were instructed to shoot through a target whose width could vary from trial-to-trial. The target consisted of an aperture within a white circle (radius: 15cm), which was always centered on the starting position (25mm^2^ red square located 15cm in front of the participant’s chest). The width of the aperture could be either 0.8cm or 8cm (Fig. 1, target phase). The width of the target on the next trial was presented well in advance of target presentation (Fig. 1, cue phase). An orange 0.8cm square or a 0.8cm × 8cm rectangle was provided as a cue in order to indicate the width of the aperture for the upcoming movement. This cue was presented 3mm below the starting position and at the same time as the starting position (~1s before reach onset). This cue disappeared at reach onset. The circle that delimited the target appeared after a delay between 700 and 850ms once the hand of the participant was stabilized inside the starting position and, in the CHOI schedule, after the participant had made his/her choice about the width of the target (see below). Once the target had appeared, participants had to reach through the target in a given time interval (500-600ms, movement phase in Fig. 1) without any information about the direction of their movement (see below). As soon as the hand crossed the circle, it left an imprint in order to provide accurate feedback about the accuracy of the movement (Fig. 1, feedback phase). In addition, the color of this imprint provided feedback about the speed of the movement (blue=too slow, yellow=too fast, green=good speed). When movement duration was between 500 and 600ms, participants earned one point. They also earned an additional point if the imprint was within the aperture.

Movement onset and offset were detected online with position and velocity criteria. Movement onset was flagged when the hand left the area of the starting position and hand velocity was higher than 0.02m/s. Movement end was detected when the distance travelled was larger than 15cm. A few hundreds of milliseconds after movement end, the hand was pushed back towards the starting position while the hand cursor was removed from the screen.

The experiment was divided into blocks of trials that were separated by a one-minute break. All participants started the experiment with a practice block. During this block, they performed tens of movements (40 trials in experiment 1, 60 trials in experiments 2 and 3) towards the 0.8 cm aperture with a small hand cursor (cursor was 9mm^2^) that provided online visual feedback of hand position to the participants. This block aimed at training participants to reach to the target at the good speed. After the practice block and for the rest of the experiment, a modified hand cursor was used during the movements towards the target (Movement phase, Fig. 1). This cursor was an expanding circle that indicated the distance traveled by the hand but not the direction taken by the participant. Outside the movement period, the hand cursor was a 9mm^2^ square. When the hand exited the circle, the small cursor indicated again the actual position of the hand. In addition, an imprint that was located on the circle indicated exactly where the hand of the participants crossed the boundary of the circles in order to provide accurate endpoint performance.

For 20% of the trials of all experimental blocks (i.e. not during the practice blocks), the hand path was constrained by stiff virtual walls (perturbation trials). These walls (Scheidt et al., 2000) were created by applying a stiff unidimensional spring (spring stiffness: 2500 N/m and viscosity: 25 Ns/m) and acted as a mechanical guide that directed the hand towards the center of the target or 2cm either on the left or the right of it. The lateral deviation observed during these trials was less than 0.5 mm In these trials, the actual position of the hand was concealed to the participant by 1) using the circular feedback during the movements 2) displaying the feedback imprint at the center of the target when the subjects crossed the target and 3) showing the projection of the actual position of the hand on the midline rather than its actual position once the hand exited the circle. To make sure that our results were not influence by this fake feedback, we removed all the trials immediately following a perturbation trial from all analyses. These trials were randomly interspersed throughout the experimental blocks.

In experiment 1, we tested the ability of the participants to voluntarily modulate their behavior when they themselves select the target width. This experiment started with a block of at least 120 trials during which participants selected the width of the target on each trial (CHOI schedule). The actual number of trials in this first block varied from participant to participant as the block was stopped only once the participant had chosen 60 times each of the two targets. Before the appearance of the starting position, participants indicated orally to the experimenter the width of the target that they would like to receive on the next trial. The operator then selected the correct target size for the upcoming movement. This first block was followed by two blocks of 90 trials each. In the first block (RND schedule), target width was varied pseudo-randomly (the order of trials was prearranged before the experiment). In the second block (BLK schedule), half of the participants experienced the narrow target for 45 trials followed by another 45 trials with the wide target. The order of target width was counterbalanced for the other half of the participants. In summary, experiment 1 consisted of three different schedules: CHOI, RND and BLK.

In experiment 2, we tested the ability of the participant to form a memory of the optimal behavior exhibited during the BLK schedule in a later random schedule. In this experiment, during the first and second experimental blocks (66 and 90 trials, random schedule, RND1), the target width was varied randomly (i.e., for block 2, the order of trials was prearranged before the experiment,) from trial to trial. During the third one (blocked schedule, BLK), half of the participants experienced the narrow target for 60 trials followed by another 60 trials with the wide target. This order was counterbalanced for the other half of the participants. During the fourth block (RND2, 120 trials), the width of the target was again pseudo-randomly presented from trial to trial. In summary, experiment 2 consisted of three schedules: RND1, BLK and RND2.

In experiment 3, we tested the ability of the participants to update their motor program from one trial to the next. This experiment consisted of four different blocks. During the first and second experimental blocks (66 and 90 trials, random schedule, RND), the target width was varied randomly from trial to trial. During the third one (Fig. 1C, mixed schedule, MIX), half of the participants experienced 60 trials where the narrow target was presented in 75% of the trials (high probability trials) and the other 25% (low probability trials) were randomly interspersed trials with the wide target. These 60 trials were followed by another 60 trials with high probability trials with the wide target (75% of the trials) and low probability trials with the narrow target). The order of target width associated with high-probability trials was counterbalanced for the other half of the participants. In other words, all the participants experienced high and low probability trials with both the narrow and wide targets. Importantly, 8 of the 15 low probability trials were perturbation trials. During the fourth experiment block (BLK), the width of the target was blocked. In summary, experiment 3 consisted of three schedules: RND, MIX and BLK.

### Data analysis

In perturbation trials, we used the signed lateral force exerted by the participants against the wall of the channel as a proxy of their willingness to go towards the center of the target. Force measures were low-pass filtered (second order Butterworth filter with cutoff: 75Hz). The average lateral force from straight ahead perturbation trials was subtracted from the measures of lateral force during rightward or leftward perturbation trials separately for each schedule and target width separately. These corrected measures were sign reversed for the rightward perturbations such that a larger positive force represents a larger reaction to the perturbation. To quantify the reaction to the perturbation, we extracted the lateral force exerted by the participants when they were 2cm away from the target. These 2cm allowed us to avoid late correction of the movements or the period where the cursor was back to normal size. To assess the effect of time on this variable, the same measure was also taken at 7cm (~240ms into the movement).

In unperturbed trials, our proxy for movement performance was the lateral position of the hand when it was 2cm away from the target. This measure was considered as movement endpoint. This measure of movement endpoint was highly correlated with the movement endpoint when the hand crossed the target (R>0.8) but had the advantage to contain many more points. Indeed, participants tended to stop their movement before reaching the target (percentage of such movement varies between participants and experiments). In such case, the value at 15cm cannot be computed. Unperturbed trials were excluded from these analyses if they immediately followed a perturbed trial. To assess how the performance of one trial affected the performance on the next trial, we computed lag 1 autocorrelation of endpoint errors between pairs of trials. Note that pairs of consecutive trials that include a perturbation trial were also excluded from this analysis.

In experiment 1: ANOVA was used with schedule (CHOI, RND, BLK) and target width (narrow and wide) as within-subjects factors.

In experiment 2: ANOVA was used with schedule (RND1, BLK, RND2) and target width (narrow and wide) as within-subjects factors.

In experiment 3: ANOVA was used with schedule (RND, MIX-high probability, MIX-low probability, BLK) and target width (narrow and wide) as within-subjects factors. The factor MIX-high corresponded to the 75% of trials that had the same target width while the factor MIX-low represented the remaining 25%.

In the cognitive task switching experiment, the reaction time was taken as the time between the appearance of the symbol on the screen and the force magnitude (in any direction) reached a threshold of 6N (which required a strong response from the participant). Then we computed the median reaction time of the correct trials for each epoch separately.

To assess how much the performance on one trial was taken into account in the next trial, we followed the method proposed by Van Beers (van Beers, 2009; van Beers et al., 2013a, 2013b) who reasoned that correlation between errors on consecutive trials should be low in the presence of trial-to-trial error correction mechanisms while it should be higher if error correction mechanisms were absent (random walk). To assess this correlation, we took into account all the pairs of consecutive trials that did not contain a perturbation trial. For each of these pairs, we considered the lateral position of the hand shortly before reaching the target (see above). Then we computed the correlation coefficient between the lateral positions in these pairs of trials.

In order to take into account the different levels of force observed in the different experiments (e.g. due to reaching or shooting), the analysis of the switching cost across the experiments from this paper and from the previous one (Orban de Xivry, 2013) was carried on the normalized lateral force obtained by dividing the lateral force by the average force.

## Results

### Self-selection of target width does not alleviate the inability to switch between two different optimal control policies

In the first experiment, participants were asked to randomly select the width of the target before each trial (CHOI schedule) in order to test whether self-selection of target width would allow the participants to optimize their reaching movements on a trial-to-trial basis during a random schedule. This block was then followed by two other blocks. In the first one, the target width was randomly changed from trial-to-trial (random schedule, RND) while in the second, the target width remained constant for 60 trials before switching to the other target width (block schedule, BLK). In each of these three schedules, we analyzed how participants responded to perturbations during their movements in function of the target width. By doing so, we make the assumption that participants would use the same control policy in perturbed and unperturbed trials because the perturbation trials were unpredictable. That is, the response to the perturbation reflects the control policy that is used in unperturbed trials. In addition, most of the participants did not actually realize that in some trials, their hand was perturbed away from its trajectory.

Both in the CHOI and RND schedules, participants slightly modulated the force that they exerted against the perturbation with target width. However, the modulation of the force with target width was significantly smaller than what was observed in the BLK schedule.

**Figure 2.**
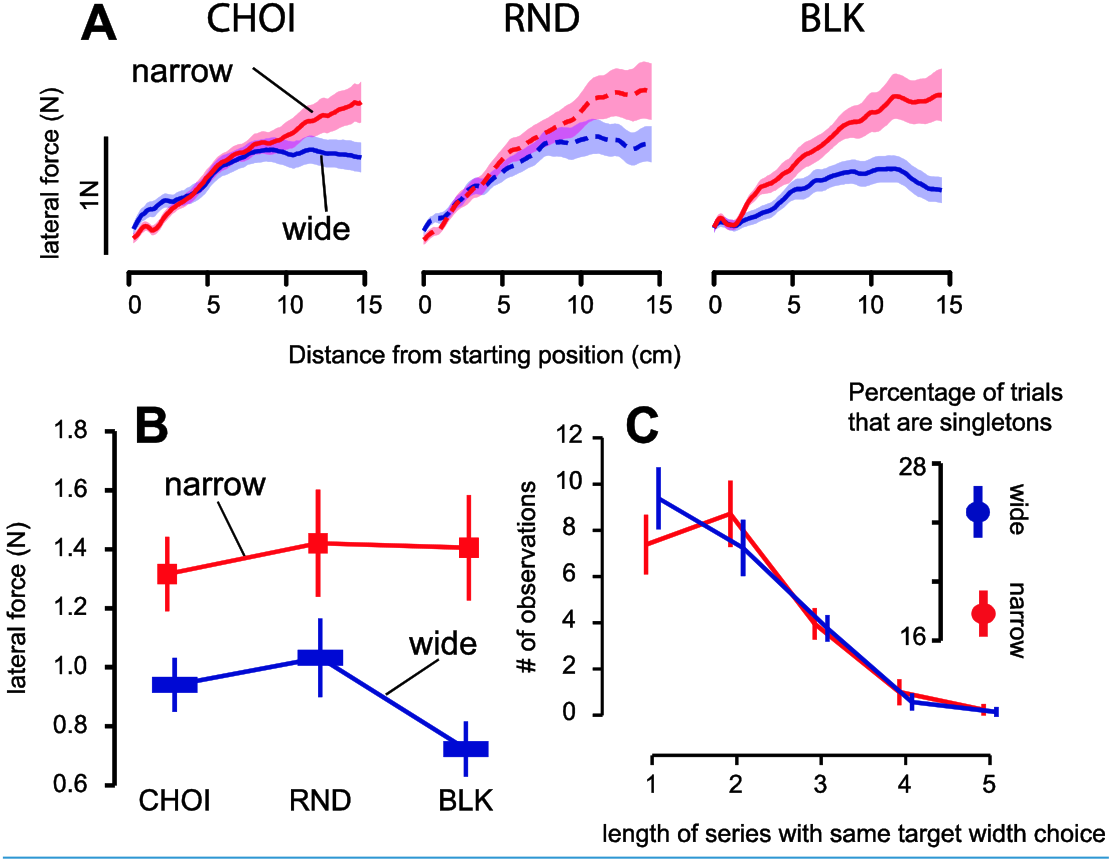
Self-selection of target width does not improve optimality of behavior in comparison with the random schedule. A) Average force profiles across all participants recorded during the perturbation trials for each schedule (CHOI: self-selection of target width; RND: random schedule; BLK: blocked schedule) and each target width separately (red: narrow target; blue: wide target). In these plots, the forces are represented against the distance from the starting position in order to match the level of perturbation. Shaded area around each curve represents standard error of the mean. B) Average force (N) recorded at 13 cm in the force profiles in function of target width and schedule. Error bars are standard error of the mean. C) Average distribution of the length of series of trials with the same target width (i.e. how many trials in a row had the same target width). Error bars are standard error of the mean. Inset in panel C) Proportion of trials where the target width is different from the target width on the previous and next trials.

This observation was confirmed by a repeated-measure ANOVA with schedule (CHOI, RND and BLK) and target width (narrow or wide) as within-subject factors. The ANOVA yielded an interaction between target width and schedule (F(2,40)=6.9, p=0.0026). Further inspection of this interaction revealed that the difference in force for the narrow and wide targets (ΔF) was larger in the BLK schedule than in the two other schedules (paired t-test: ΔF_BLK_ vs ΔF_CHOI_: t(20)=-3.12, p=0.005; ΔF_BLK_ vs ΔF_RND_: t(20)=-3.04, p=0.006). There was no statistical difference in ΔF between the CHOI and RND schedules (t(20)=-0.19, p=0.84). The modulation of force with target width was nevertheless significant both in the CHOI and in the RND schedules (ΔF >0: CHOI: t(20)=5.3, p<0.001; RND: t(20)=4.2, p<0.001). Overall, these results suggest that despite being able to select the target width on each trial, there was only a limited modulation of the response to the perturbation with target width during the CHOI schedule compared to the BLK schedule.

In the CHOI schedule, participants were explicitly instructed to select the target width randomly before each trial. However, they were not able to do so. Most of the participants tended to select the narrow target for several trials in a row while they more often selected the wide target for a single trial before switching back to the narrow target. Therefore, after 80 trials, most participants had performed more trials with the narrow target (on average, ~42 trials) than with the wide target (on average ~38 trials). In addition, the proportion of single trials with the narrow and wide targets differed (narrow vs wide: 18% vs 25%, t(20)=-4.25, p=0.0003). That is, there were less isolated trials (hence, more trials in a row) for the narrow target than for the wide target. If the participants wanted to minimize effort, they should have a bias towards the wide target (i.e. they should have chosen the wide target more often than the narrow target).

### Relationship between switching cost for motor and cognitive tasks

The subjects who participated in the first experiment also participated in a cognitive task switching. In this task, participants had to indicate either the shape or the color of a symbol that appeared in the middle of the robot workspace (Fig. 1A). Subjects were able to learn the rules in a satisfactory way (RND: 85% accuracy; BLK: 90% accuracy). When the task instruction changed randomly from trial to trial, the reaction time of the participants was slightly above 1s (1085±47ms, mean ± SE). In contrast, the reaction time when the task instruction remained unchanged for several trials in a row was 400ms shorter (640±45ms, mean ± SE, t(20)=13.66, p<0.0001). These 400ms represent the switching cost between the RND and BLK schedules (Fig. 3A). Similarly, the switching cost in the motor task was around 0.3N, which represents the difference in modulation of the force with target width between the RND and BLK schedules (Fig. 3C). Interestingly, we found that the cognitive and motor switching costs were correlated across participants (Fig. 3B, r=0.57, t(20)=3.06, p=0.007). That is, participants for whom the schedule had a small effect in the cognitive task (reaction time in RND not much worse than in BLK) were better able to modulate their force in the RND schedule in comparison to the BLK schedule. This relationship suggests that both motor and cognitive switching cost might stem from a general mechanism of the central nervous system.

**Figure 3.**
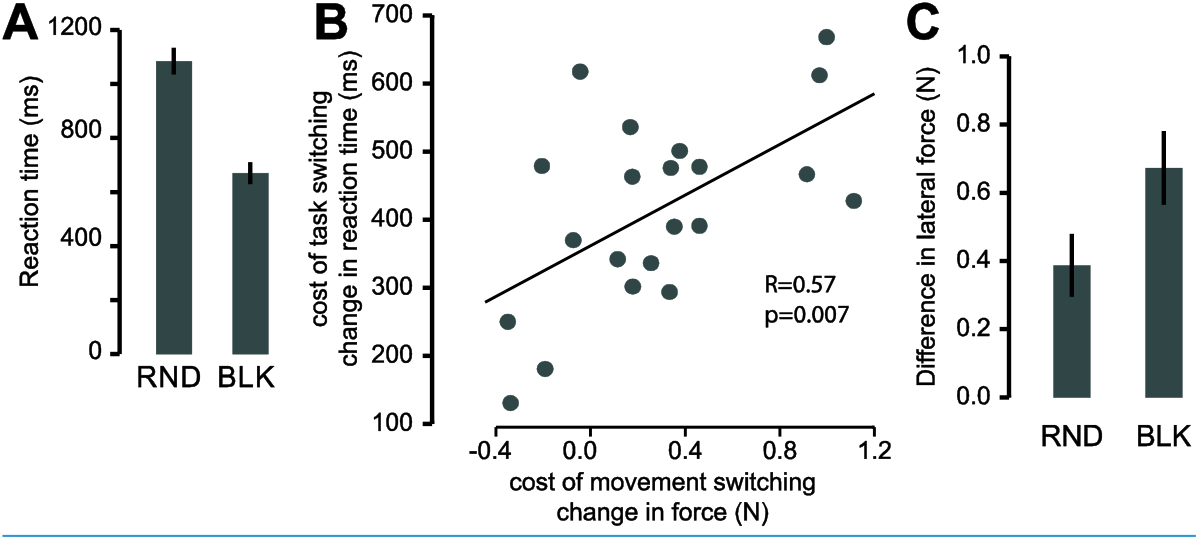
Link between cognitive and motor switching cost. A) Reaction time (ms) in the random (RND) and blocked (BLK) schedules during the cognitive task switching. The difference between the two bars (RND-BLK) represents the average cognitive switching cost (ms). B) Correlation between the cognitive and motor switching costs. C) Amount of modulation in the force exerted against the perturbation with target width (force for narrow target minus force for wide target) for the RND and BLK schedules. The difference between the two bars (BLK-RND) represents the motor switching cost (N).

### Knowing the optimal motor program does not help behaving optimally in the random schedule

The results from these experiments and previous studies on optimality of motor behaviors (de Rugy et al., 2012; Kistemaker et al., 2014) are actually confounded by the possibility that participants might not know what the optimal behavior is because they have never used it. Therefore, in the second experiment, we tested whether the absence of optimality during the random schedule was due to the inability to adopt the optimal motor behavior or to the ignorance of the optimal motor behavior. Indeed, so far in all the experiments, participants had always experienced the RND schedule before the BLK schedule. Again, we found in the first random schedule that participants applied more force against a perturbation when the target was narrow than when it was wide (paired t-test, t(19)=2.37, p=0.028). However, this difference between the force applied against the perturbation for the narrow and wide targets was much larger during the blocked schedule (interaction between schedule (RND1 and BLK) and target width, F(1,19)=7.08, p=0.015). We then tested whether the participants were able to use the learned optimal behavior in a second random schedule (RND2). We found that, when participants experienced the random schedule again, the difference in force applied against the narrow and wide targets was again smaller than the modulation of force with target width observed during the blocked schedule (interaction between schedule – BLK and RND2 - and target width, F(1,19)=7.09, p=0.015) and they did not differ significantly from the force in the first random schedule (interaction between schedule – RND1 vs. RND2 - and target width: F(1,19)=0.25, p=0.62). These data suggest that even though participants have used the optimal motor behavior in the preceding blocked schedule, they were unable to use it during the following random schedule. In other words, learning and using the optimal control policy in the blocked schedule does not lead to a better performance in the subsequent random schedule.

**Figure 4.**
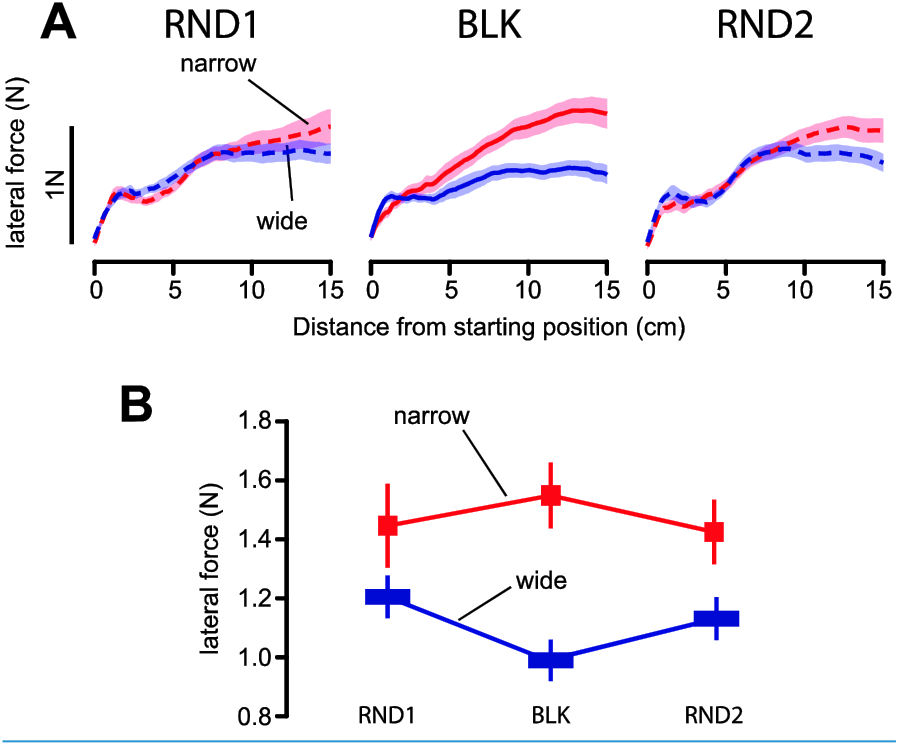
Knowing the optimal motor program does not improve behavior in a subsequent random schedule A) Average force profiles across all participants recorded during the perturbation trials for each schedule (RND1 and RND2: random schedule; BLK: blocked schedule) and each target width separately (red: narrow target; blue: wide target). In these plots, the forces are represented against the distance from the starting position in order to match the level of perturbation. Shaded area around each curve represents standard error of the mean. B) Average force (N) recorded at 13 cm in the force profiles in function of target width and schedule. Error bars are standard error of the mean.

### Inability to instantaneously switch between motor programs

While the previous experiment aimed at emphasizing the inability to switch between motor behaviors despite knowing the optimal motor program, we wanted next to identify the cost of switching when target width changed unexpectedly during a blocked schedule. To do so, twenty other participants experienced a new schedule (MIX schedule) where 75% of the trials had a given target width (MIX-high: high probability trials, e.g. narrow target) and 25% (MIX-low: low probability trials) had the other target width (e.g. wide target). Half of these low probability trials were perturbation trials.

**Figure 5.**
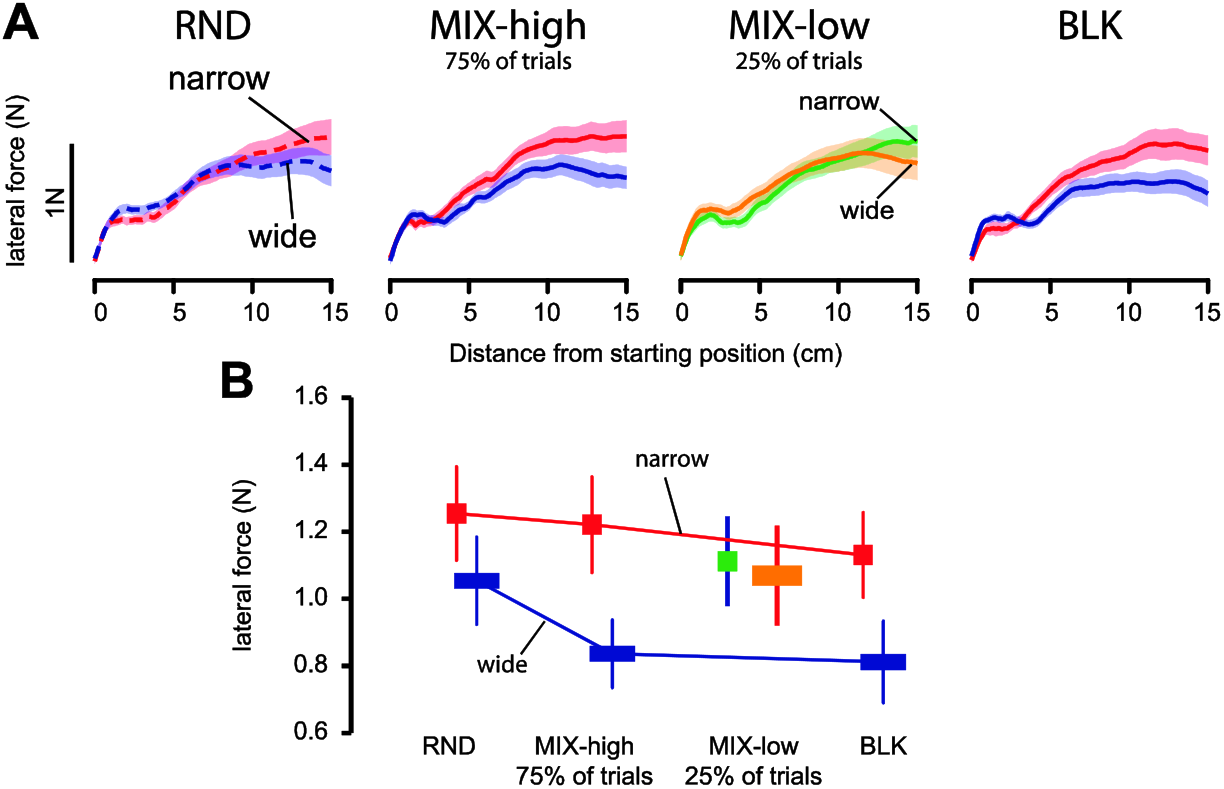
Change in motor program over one trial A) Average force profiles across all participants recorded during the perturbation trials for each schedule (RND: random schedule; MIX-high: high probability trials in the mixed schedule; MIX-low: low probability trials in the mixed schedule; BLK: blocked schedule) and each target width separately (red: narrow target; blue: wide target). In these plots, the forces are represented against the distance from the starting position in order to match the level of perturbation. Shaded area around each curve represents standard error of the mean. B) Average force (N) recorded at 13 cm in the force profiles in function of target width and schedule. Error bars are standard error of the mean.

In this group, difference in force profiles were again more marked during the blocked schedule than during the random schedule. In addition, during the MIX schedule, the force that the participants exerted against the perturbation was clearly modulated by target width for the most frequent target width but not for the target width presented in the low-probability trials. These observations were confirmed by an interaction between target width and trial type (random, MIX-high, MIX-low and BLK: F(3,57)=2.9267, p=0.041). Post-hoc Tukey t-tests revealed that target width modulated force only for the high probability trials in the MIX schedule and for the blocked schedule (p=0.001= and p=0.013) but neither in the random schedule nor in the low probability trials (p=0.31 and p=0.99). In other words, the behavior in the low probability trials was close to what was observed in the random schedule.

### Additional knowledge from our large pool of subjects (N=111)

In the three experiments presented here and the four ones presented in a previous paper (N=111 subjects), we found a very strong effect of schedules on the ability to modulate the response to a perturbation (Fig. 6.A; interaction between target size and schedule (RND and BLK): F(1,110)=35.18, p<0.00001, partial eta square: 0.24). In order to take into account the different levels of force observed in the different experiments (e.g. due to reaching or shooting), this analysis was carried on the normalized lateral force (obtained by dividing the lateral force by the average force).

In addition, this database allows us to have a look on the actual difference between RND and BLK. Indeed, there appears to be quite some variability across experiments (Fig. 6.B). While in some experiments (e.g. #1), the increase in force for the narrow target from RND to BLK was pronounced, it was absent or small in other experiments (#6 and #7). Similarly, for the wide targets, there was an important decrease in force from the RND to the BLK schedule is some experiments (e.g experiment #7) but not in others (e.g. #1). On average, on the basis of the 111 subjects, we can conclude that the schedule modulated the force both for the narrow target (RND vs. BLK: Tukey Post-Hoc: p=0.0002) and for the wide target (Tukey post-hoc: p=0.0001). This observation allows us to reject the possibility that subjects were lazy and pushing too hard for the wide target as a safety measure. In addition, we observe that the same control policy was not used for both narrow and wide target in the RND condition given that the response to the perturbation was modulated by target width (Tukey Post-Hoc: narrow vs wide target in RND condition: p=0.0001). However, this modulation was smaller in the RND condition than in the BLK condition as indicated by the significant interaction.

**Figure 6.**
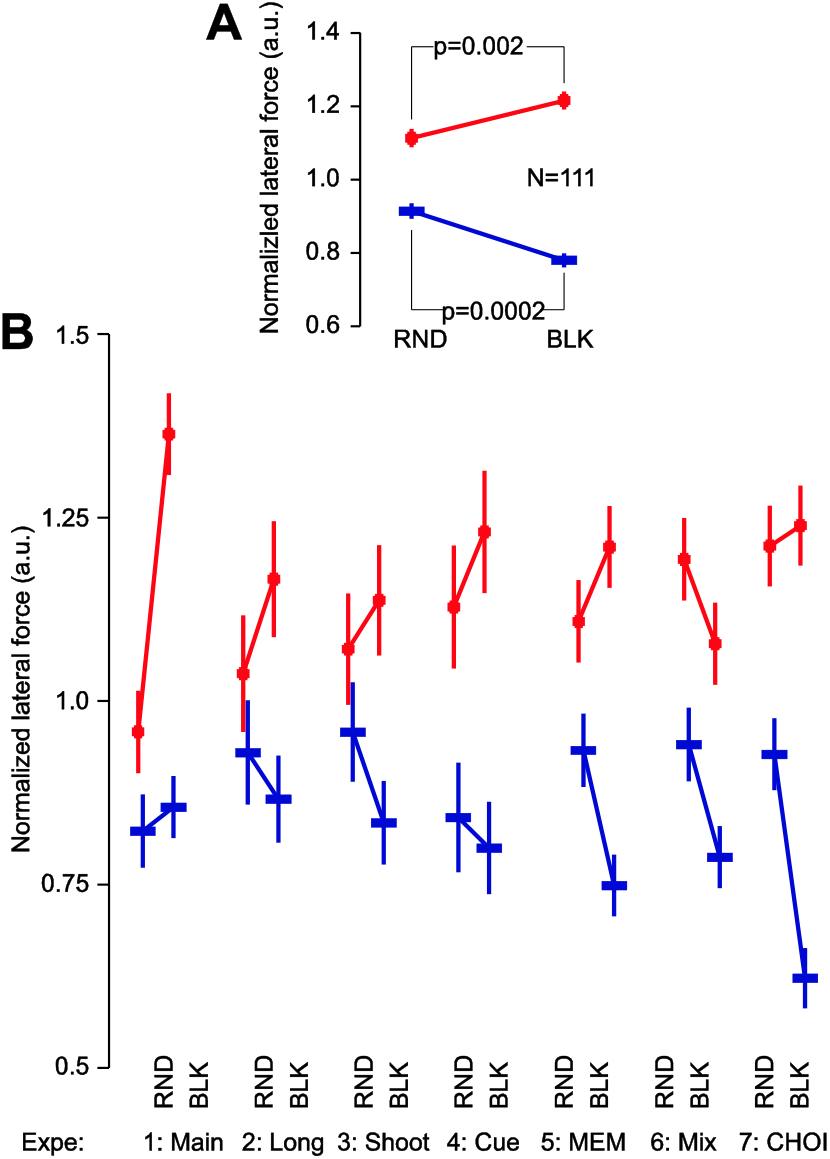
Analysis of the switching cost for the experiments reported in this paper (1 to 3) and in our previous (4: Main; 5: Long; 6: Shoot; 7: Cue experiments). Panel A: Normalized lateral force for the two schedules and target widths for the 111 subjects. B: Representation of the variability of the observed effect across the seven experiments. Error bars represent standard error of the mean.

Given the nature of the perturbation (position-dependent perturbation), it is difficult to judge how early the motor system responded differently to a narrow or wide target. Nonetheless, an ANOVA on force measures at a distance of 7cm (around 240ms after movement onset) from the starting point revealed the same target width × schedule interaction (F(1,110)=37.8, p<00001).

### Absence of history effect in response to the perturbation

Costs in cognitive task switching are highlighted both by the difference between random and blocked schedules but also by a difference in performance between switch trials (previous trial had a different goal from the current one) and non-switch trials (previous trial was similar to the previous one).

**Figure 7.**
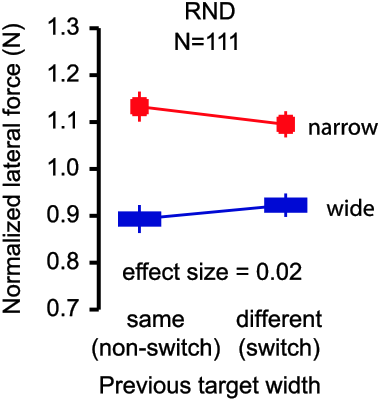
History effect: influence of the target size from the previous trial on the normalized lateral force (N) exerted on the current trial. This graph summarizes the data from 111 participants (61 from this paper and 40 from Orban de Xivry 2013). Data from the RND schedules only. Error bars represent standard error of the mean.

In contrast to cognitive switching cost, we did not find differences in performance between switch (change in target width between the previous trial and the current one) and non-switch trials (same target width on the previous trial as on the current one). That is, we expected that the force to a perturbation would be larger on one trial when it was preceded by a trial with the narrow target and smaller when it was preceded by a trial with the wide target. In other words, we expected an interaction between target width and switching state (switch vs non-switch trials). In our large pool of participants (N=111, Fig. 7), we found that the response to the perturbation was very weakly influenced by the switching (partial eta square: 0.02; F(1,109)=2.06, p=0.15). As always, the absence of significance could be due to an absence of power. For this reason, we reported the effect size of the effect: 0.02, which suggests that, even if the effect exist, it should be considered as a small effect (Cohen, 1988). In addition, such history-dependent effects, in other contexts, are typically observable with much smaller sample size (2 in Kowler et al., 1984; 8 in Witney et al., 2001; 10 in Franklin et al., 2008; 7 in Tabata et al., 2008). This suggests that the smaller modulation of the response to perturbation with target width during the random schedule is not due to the inability for the participants to converge to the optimal solution. That is, it is not a gradual process where participants adjusted their movement planning trial after trial.

### Trial-to-trial changes in the motor program

**Figure 8.**
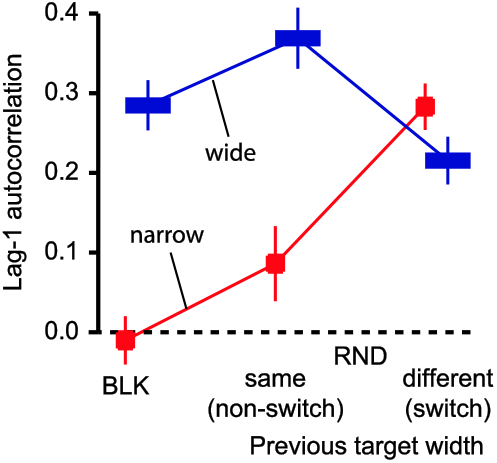
Trial-to-trial changes in movement endpoint: Lag-1 autocorrelation of movement endpoint error for different pairs of trials from the RND and BLK schedules. Error bars are standard error of the mean. This graph summarizes the data from 111 participants (61 from this paper and 40 from Orban de Xivry 2013).

Updating the planned direction of the next movement as a function of the error experienced in the previous trial leads to low lag-1 autocorrelation between consecutive movement endpoint errors (van Beers et al., 2013a). In contrast, a high lag-1 autocorrelation is associated with little or no trial-to-trial changes in motor performance (similar to a random walk). In the three experiments presented here and the four presented in a previous paper (N=111 participants), we found that, during the BLK schedule, the lag-1 autocorrelation was almost zero for the narrow target but was much larger for the wide target (Fig. 8; N=111, −0,034±0.027 vs. 0.24±0.029, t(110)=-8.12, p<0.0001, partial eta square = 0.37). We also looked at the lag-1 autocorrelation of endpoint error between pairs of trials with or without a change in target width. Interestingly, during the random schedule, pairs of consecutive trials with narrow targets or with wide targets exhibited the same behavior as in the BLK schedule (smaller for consecutive trials with the narrow target than for consecutive trials with the wide target, 0.11±0.04 vs. 0.36±0.033, t(110)=-5.8, p<0.0001, partial eta square = 0.24). In contrast, a change in target width appeared to reduce the ability to take the performance from the previous trial into account as lag-1 autocorrelation between pairs of trials where the target width changed was high in both cases (narrow to wide: 0.2±0.025; wide to narrow: 0.29±0.025). Overall we found an interaction between switch and target width variables (Fig. 8, F(1,110)=47.6, p<0.0001). Post-Hoc test suggests that the lag-1 autocorrelation was lower for the pair of trials with narrow targets than for the three other pairs (Dunnett’s t-test: SS vs SL: p<0.001; SS vs LS: p= 0.023; SS vs LL: p<0.001). This pattern appears similar for the two datasets (2013 vs. 2015; interaction between switch, target width and dataset: F(1,109)=.04468, p=0.83).

In the BLK schedule, repetitiveness did not disengage the participants as they were more accurate and more often inside the target in the blocked condition (46% of the time) than in the random condition for the small target (40% of the time). Therefore, the success rate increased by 15% for the small target between the RND and BLK conditions (t(110)=4.51, p=0.00002). So, even though the RND schedule was potentially more engaging than the BLK schedule, participants were less successful during that schedule.

Finally, we did not notice any cost of switching in the median reaction time as confirmed by an ANOVA on the median reaction time with schedule and target width as within-subject factors (main effect of target width: F(1,110)=1.44, p=0.23, partial eta square = 0.012; interaction: F(1,110)=1.79, p=0.18).

## Discussion

This study makes two important points about movement planning. First, the inability of our participants to behave optimally in a changing environment suggests the presence of a switching cost. More specifically, in our experiments, participants exhibited different motor behaviors in the random and blocked schedules, even when the participants themselves selected the width of the target that they will reach toward or when the participants have previously demonstrated the knowledge of a better motor program for the task at hand. Second, participants were unable to use the endpoint error on one trial in order to adjust their planning of the next movement if the target width changed. These results directly speak over recent theories of motor planning (Wong et al., 2015), which suggest that movement initiation results from a series of processes that can be separated into two categories. Processes from the first category determine the motor goal (the ‘what’ pathway: target extraction and selection, attention, etc.) and are largely responsible for variation in reaction time while the processes of the second category are responsible for action selection and for the specification of the movement kinematics (the ‘how’ pathway). Given the absence of reaction time modulation in our task, we believe that our results speak directly to the limitations of the ‘how’ pathway, which encompasses both action selection and movement specification.

### Reduced ability to learn from the error in the previous trial in order to adjust endpoint position

In simple reaching movements like in this study, action selection refers to the selection of the next aiming direction, which varies from trial-to-trial. Such trial-to-trial change in aiming direction has been considered as a learning from error process where the aiming direction on a trial is equal to the aiming direction on the previous trial plus a fraction of the error (van Beers, 2009). Such a process causes the correlation of endpoint errors between two consecutive trials to be small if the errors are taken into account (i.e. for the narrow target) and a positive auto-correlation when it is not (i.e. for a wide target, where errors do not matter) (van Beers et al., 2013a). However, this theory is inconsistent with the auto-correlations observed during the random schedule. When the target presented on one trial was narrow, participants updated the aiming direction for their next trial only if the target on the next trial was also narrow but not if it was wide (Fig. 8). This suggests that the aiming direction cannot be updated before the target width of the next trial is known and that participants held the error in memory until they saw the target width of the next trial. This scheme appears very unlikely. If this scenario was possible, then the error observed in trials with wide target should be used if the next target was narrow. To the contrary, the success rate in these trials with narrow target was low compared to success rate in the BLK schedule. Alternatively, it is possible that the subjects were unable to estimate the size of the error with respect to the center of the target because its ends were too far away. We believe that this explanation does not hold as the wide target is 8cm wide (~9 deg of visual angle) and people can accurately estimate the middle of a 20cm line in a line bisection task (error of 1% reported in Milner et al., 1992) or can accurately maintain their eye at the center of large targets (30 deg of visual angle) defined only by illusory contours (Wyatt et al., 1994). In summary, if the theory of error-based learning as suggested by Van Beers is correct, it may only apply to repetitive movement to a single target, which represents a very limited situation in everyday life, except maybe in dart throwing (van Beers et al., 2013b).

This suggests that an alternative framework for the update of the aiming direction is necessary. This framework should have the same characteristics as the Monte-Carlo Markov Chain framework (MCMC) proposed by Haith and Krakauer (2014). In this framework, the update of the aiming direction does not depend on the error but on the rewarding consequences of a trial (Skinner, 1981), which depends on target width as is the case for optimal control (Nashed et al., 2012). That is, the previous aiming direction is replaced by the current one with some probability that depends on the value of the previous and current aiming direction (Haith and Krakauer, 2014). However, the update of the aiming direction can only take place when the cost function is unchanged between two consecutive trials as one cannot compare values from different scales (like comparing apples and oranges). This would also explain why participants tended to repeat consecutive trials with a narrow target (Fig. 2C) when asked to select the target width on each trial. Choosing the narrow target repeatedly allowed them to update their aiming direction in function of their past performance. To summarize, the MCMC framework provides a convincing explanation for the absence of updating of movement plans when target width changes and for the repetitive choice of the same target width when the target is narrow.

### Inability to be optimal

After action selection, the kinematics of the movement need to be specified (Wong et al., 2015). Our data suggest that the optimality of the control policy was disrupted by the frequent changes in target width. Optimal control predicts that every movement should be optimal (Todorov and Jordan, 2002; Nashed et al., 2012), independently of the schedule. However, we found that the modulation of the response to perturbations with target width was much reduced but well present in the random schedule compared to the blocked schedule.

This inability to modulate the feedback control policy adequately in the random schedule led to a cost in accuracy and a cost in effort. That is, participants were unable to reach the same accuracy level for the narrow target in the random schedule as in the blocked schedule. In addition, participants spent too much energy controlling the hand in the random schedule when the target was wide compared to in the blocked schedule. This decrease in performance due to the randomization of condition is a general finding that extends far beyond our simple task (Elliott and Allard, 1985; Edin et al., 1992; Horak and Diener, 1994; Khan et al., 2002; Pruszynski et al., 2008; Afsanepurak et al., 2012). Here, we provide for the first time, an account for this phenomenon.

This absence of optimality in the random schedule is not due to the long time required to reach the optimality. Indeed, we observed an abrupt change in force response after the (unique) change in target width in the blocked schedule (Orban de Xivry, 2013). Second, optimization of the control policy did not lead to trial-to-trial changes in force response against the perturbation. Indeed, there was no effect of the target width experienced on the previous trial on the force applied on the next trial (Fig. 7). Therefore, this suggests that our effect is not due to the fact that it takes several trials in order to reach the optimal behavior. In contrast, in the blocked schedule, optimality was achieved in the first few trials (Orban de Xivry, 2013). It also rejects the idea that performance in the random schedule is deteriorated due to a history-dependent effect. In contrast, such history-dependent effects are observed in predictive eye movements (Kowler et al., 1984; Tabata et al., 2008) for which the memory of target trajectory is updated after each trial (Orban de Xivry et al., 2013) and for grasping movements where an internal model of the object weight is built trial after trial (Johansson and Westling, 1988; Loh et al., 2010). Finally, our data goes against the possibility that a single habitual control policy was used in the random schedule because we observed a significant modulation of the behavior with target width in the random schedule as well. However, this modulation was smaller than in the blocked schedule.

In many complicated tasks, optimality might be difficult to reach (Kistemaker et al., 2010, 2014; de Rugy et al., 2012). In addition, it is actually unknown whether participants would have been able to execute the optimal motor behavior because they had never performed optimal movements in these contexts before. In contrast, in our task, we know, thanks to the behavior in the blocked schedule, that participants could select a better feedback control policy but that they did not. Indeed, in experiment 2, despite having used a better (more accuracy for the narrow target and less energy wasted for the wide target) feedback control policy in the blocked schedule, participants were unable to use these control policies in the subsequent random schedule. This observation supports our claim that movements in the random schedule were not optimal when compared to the blocked schedule.

In other words, there has to be a cost associated with changing the feedback control policy, i.e. a switching cost. If no such cost existed, then participants could use their best (less costly) strategy to perform the task (the one used in the blocked schedule). One possibility is that the control policy is optimized in function of the context, allowing a more limited modulation of the control policy in the random schedule compared to the blocked schedule. This view is also compatible with the idea of a switching cost for motor control as this cost would be responsible for this limitation.

In the case of multiple targets, several motor programs are prepared together during the planning stage (Cisek and Kalaska, 2005, 2010) and the non-selected one is later inhibited. The current study leads to the question of whether there are multiple programs prepared when a unique target location must be associated with two different control policies (for the narrow and wide targets)? In this case, inhibition of the non-selected motor program could lead to subsequent switching cost as it became harder to retrieve the inhibited motor program (Duque et al., 2005; Duque and Ivry, 2009; Mars et al., 2009) or the inhibited cognitive task (Mayr et al., 2000, 2006). Alternatively, the switching cost might arise from the fact that a new control policy must be selected each time the target width changes.

To summarize, we do not question the fact that, following optimal feedback control, feedback gains are determined before movement in order to bring the hand on the target or to respond to potential perturbations during the movement. However, we want here to highlight that this optimization of the feedback control law is not as flexible as suggested by many authors and that switching between control policies carry some costs.

### How similar is the phenomenon described here to cognitive task switching

The motor switching cost identified here bears many similarities with the switching cost for cognitive tasks. However, it also differs from it in a number of ways. First, while we identified here a switching cost for the ‘how pathway’ (Wong et al., 2015), cognitive studies typically focus on a cost on the ‘what pathway’ which results in an effect on the reaction time (Rogers and Monsell, 1995). It is indeed critical to realize that several factors that directly influence the feedback control policy do not modulate the reaction time and that movement preparation and initiation are two independent processes (Haith et al., 2016). For instance, the influence of target size on reaction time is very limited (Quinn et al., 1980). It is believed that the feedback control policy is only selected among several possibilities. Such process does not largely influence the reaction time.

Despite this difference, self-selection of target (experiment 1) or selection of cognitive task rule (Arrington and Logan, 2004) does not abolish the switching cost. Second, while a switching cost can be identified on a single trial basis in cognitive tasks (reaction time is different in switch and non-switch trials), we failed to identify such effect in our data (Fig. 7). However, we found that the switch prevented participants from taking the performance of the previous trial into account in order to update their movement plan for the next trial (Fig. 8).

Finally, we found that motor and cognitive switching costs were correlated across participants. This correlation might stem from the fact that updating the association between a stimulus and a motor program might be controlled by a mechanism that is independent of the rationale behind the update (i.e. change in task rule or change in target width) such as the one taking place in the premotor and motor areas for cognitive and motor tasks. Cells from the cingulate cortex are differentially modulated when the movement plan needs to be updated or when it needs to be repeated (Shima and Tanji, 1998; Procyk et al., 2000). Lesions of the same area impair performance at a cognitive switching task in monkeys (Rushworth et al., 2003). The pre-supplementary and supplementary motor areas (Nachev et al., 2008) appear to also play an important role in the cognitive and motor switching processes. Cells of the pre-supplementary and supplementary motor areas only responded before switch trials and not before non-switch trials during a reaching task to two targets (Matsuzaka and Tanji, 1996). Similarly, pre-SMA first inhibits the previous motor program and then boosts the desired saccade when instructed to switch (Isoda and Hikosaka, 2007; Hikosaka and Isoda, 2010).

## Conclusion

A simple manipulation of target width during a reach experiment reveals the limitations of the mechanisms of motor planning. These data reveal that action selection might proceed via a selection by consequence mechanism and that movement specification suffers from a switching cost for motor planning. This switching cost for motor tasks shares many similarities with the switching cost identified in cognitive tasks.

## Acknowledgements

This work was supported by the Belgian Program on Interuniversity Attraction Poles and PRODEX initiated by the Belgian Federal Science Policy Office, Actions de Recherche Concertée (French community, Belgium) and the European Space Agency (ESA) of the European Union. JJO was supported by the Brains Back to Brussels program from the Brussels Region (Belgium).

## Author contributions

J.-J.O.d.X. conception and design of research; J.-J.O.d.X. performed experiments; J.-J.O.d.X. analyzed data; J.-J.O.d.X. interpreted results of experiments; J.-J.O.d.X. prepared figures; J.-J.O.d.X. drafted manuscript; J.-J.O.d.X. and P.L. edited and revised manuscript; J.-J.O.d.X. and P.L. approved final version of manuscript

